# Direct observation of the conformational states of formin mDia1 at actin filament barbed ends and along the filament

**DOI:** 10.1101/2022.06.28.497909

**Authors:** Julien Maufront, Bérengère Guichard, Lu-Yan Cao, Aurélie Di Cicco, Antoine Jégou, Guillaume Romet-Lemonne, Aurélie Bertin

## Abstract

The fine regulation of actin polymerization is essential to control cell motility, architecture and to perform essential cellular functions. Formins are key regulators of actin filament assembly, known to processively elongate filament barbed ends and increase their polymerization rate. Based on indirect observations, different models have been proposed to describe the molecular mechanism governing the processive motion of formin FH2 domains at polymerizing barbed ends. Using electron microscopy, we directly identified two conformations of the mDia1 formin FH2 domains in interaction with the barbed ends of actin filaments. These conformations agree with the open and closed conformations of the “stair stepping” model proposed by Otomo and colleagues^1^. We observed the FH2 dimers to be in the open conformation for 79% of the data, interacting with the two terminal actin subunits of the barbed end, while they interact with three actin subunits in the closed conformation. Further, our data reveal that the open state encompasses a continuum of states where the orientation of the leading FH2 domain with respect to the filament long axis varies from 108 to 135 degrees. In addition, we identified FH2 domains encircling the core of actin filaments, providing structural information for mDia1 away from the barbed end. Based on these direct observations, we propose a model of formin in interaction with the growing filament end, as well as with the core of the filament.

## Introduction

The actin cytoskeleton is essential to ensure cell motility, to regulate cell shape, architecture, and to control membrane reshaping during cell division^2–5^. A myriad of actin-binding proteins ensures the fine regulation of the assembly and disassembly of actin filaments^6,7^. A large family of dimeric proteins called formins can track filament barbed ends to control their growth^8^. Formins play a crucial role in the generation of filaments found in stress fibers, filopodia or lamellipodia^9^.

Most formins gather several functional domains: a RhoGTPase Binding Domain (GBD), a Diaphanous Inhibitory Domain (DID), a Dimerization Domain (DD), two central Formin Homology Domains (FH1 and FH2) and a Diaphanous Autoinhibitory Domain (DAD). FH1 and FH2 are ubiquitous and are key to ensure the main functions attributed to formins. The FH2 domains dimerize in a head-to-tail fashion and encircle filament barbed ends. A “post” domain is located at the C-terminal end of each FH2 domain while a “knob”, a flexible “linker” and a “lasso” domain are located towards its N-terminus. The “lasso” domain of a given FH2 interacts with the “post” domain of its counterpart to induce FH2 dimerization. The “linker” domain is flexible and was shown to be either unstructured or to be able to adopt different secondary structures such as alpha helices or beta sheets^10,11^. Residues from both “post” and “knob” domains are engaged specifically in direct interaction with actin subunits at the barbed end^11–13^. FH1 domains are viewed as semi-flexible chains of polyproline-rich domains which can recruit one or several profilin and profilin-actin complexes and tune barbed end elongation rate^14^.

The molecular mechanism governing the tracking of the actin barbed ends by the FH2 domains remains elusive. Two antagonistic models have emerged to describe formin processivity at barbed ends, based on indirect evidence from X-ray crystallography of formin-actin complexes in non-native states^1^, from biophysical assays^15^, and from molecular dynamics computations^16^. Otomo and colleagues^1^ have generated the X-ray structure of a pseudo-actin filament (with a 180° helical twist, instead of the canonical 167°) bound to FH2 domains of yeast formin Bni1p, which are wrapped around the actin polymer in a continuous helix. In this non-native structure, two actin-binding regions (ABR) could be identified within each FH2 domain, the “post/lasso” and the “knob” domains. A two-state “stair stepping” model was proposed. In this model, hypothetical conformational reorganizations of the FH2 domains take place around this structure where the FH2 dimer alternates between a closed and an open state. The closed state is derived from the observed structure by reorienting the linkers to form an FH2 dimer. In this state, the FH2 dimer would interact with three actin subunits simultaneously, and a steric clash from the leading FH2 would block monomer addition at the filament barbed end beyond the third terminal subunit. To allow the addition of an actin monomer, the authors proposed an open state, where the FH2 dimer would interact with only the two terminal actin subunits at the barbed end, leaving the “post/lasso” actin-binding interface of the leading FH2 domain free and accessible. In this “stair stepping” two-state model, it is hypothesized that, while one FH2 domain is bound to the actin filament via its two ABRs, the other FH2 domain undergoes a dynamic translocation, from one location where its two ABRs are bound to the filament (closed state), to another location closer to the barbed end with only one ABR bound (open state). After the addition of one actin subunit, the roles of the two FH2 domains are exchanged, and formins cycle through these steps to control the elongation of filament barbed ends. This two-step model was used to fit the acceleration of elongation observed when pulling on a filament growing from a surface-attached formin^15,17^ and to account for the steps observed during formin-assisted elongation thanks to an optical trap^18^.

Paul and Pollard have proposed an alternative model, named “stepping second” because the translocation of the FH2 dimer, the “stepping”, is supposedly triggered by the incorporation of a novel actin subunit^14,19^. In that model, the FH2 domains remain in the direct vicinity of the three terminal actin subunits at the barbed end, encircling them continuously. The terminal actin subunits are in rapid equilibrium between a closed conformation, with a helical twist near 180°, and an open conformation with a more canonical 167° helical twist. These conformational changes are accompanied by FH2-actin contact modulations and by the stretching of the FH2 linkers. Upon addition of a new actin monomer, the canonical 167° helical twist would be consolidated, thereby straining durably the FH2 linkers. The translocation of one FH2 domain towards the barbed end would relax this strain. After translocation, the barbed end helical twist would again be in rapid equilibrium between 167° and 180°.

The purpose of our investigations was to discriminate between these two models, by using electron microscopy to directly observe the conformations of the formin mDia1 FH2 dimer at the filament barbed end. To achieve this, we have optimized the sample preparation to enhance the density of single short filaments (Supplementary Figure 1) and adapted the image processing workflow to the processing of our filamentous ends. Indeed, only a few reports describe the structure of proteins at either the barbed or pointed ends of actin filaments, and they involve proteins that stably cap the ends and thus neutralize the actin filament dynamics^20,21^. We report here the direct observation in EM of FH2 dimers with elongating actin filament barbed ends (figure 1). This observation allows us to identify two major conformations (figure 2 & 3) which argue in favor of the “stair stepping” model. Furthermore, we report a conformational variability of the open state (figure 3). We also resolved FH2 dimers in interaction with the core of the filament (figure 4). This indicates that formins can find themselves away from the barbed end, more frequently than previously thought.

**Figure 1.**
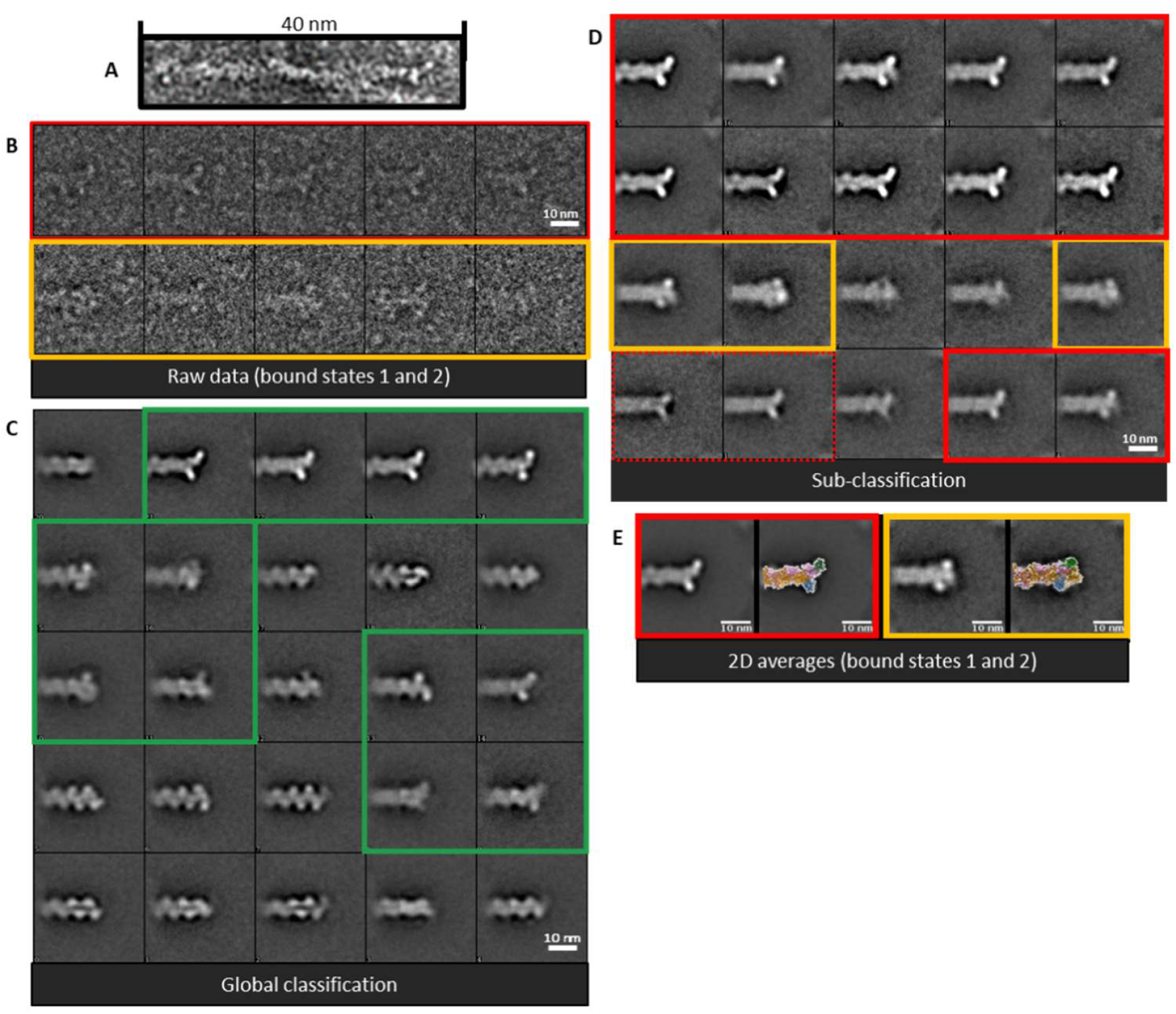
2D patterns observed at actin filament ends in the presence of formins. **A**. Image of an actin filament end displaying an extra density. **B**. Raw images corresponding to the majority (top row, *Red window*) or the minority (bottom row, *Orange window*) 2D classes of formin-bound actin filament barbed ends. **C**. SPIDER 2D classes generated from actin filament ends (20,922) observed by negative staining electron microscopy after actin filament sonication, followed by incubation with 100 nM mDia1 formins. *Green windows*: 2D classes of actin filament ends showing additional densities attributable to bound formins **D**. 2D classes generated from a subselection of actin filament ends, formin-bound candidates (**A**. *Green windows*). *Red windows*: Majority 2D classes showing additional densities protruding from the actin filament tip. *Orange windows*: Minority 2D classes showing additional densities embedded at the actin filament tip. **C**. Global class averages of the majority (*Red window*, 3,804 particles) or the minority (*Orange window*, 907 particles) configurations observed alone (left) or overlayed with the atomic models of a barbed end interacting with a formin (right) in the open (*Red window*) or in the closed (*Orange window*) states according to the “stair-stepping” model (ref Otomo).

**Figure 2.**
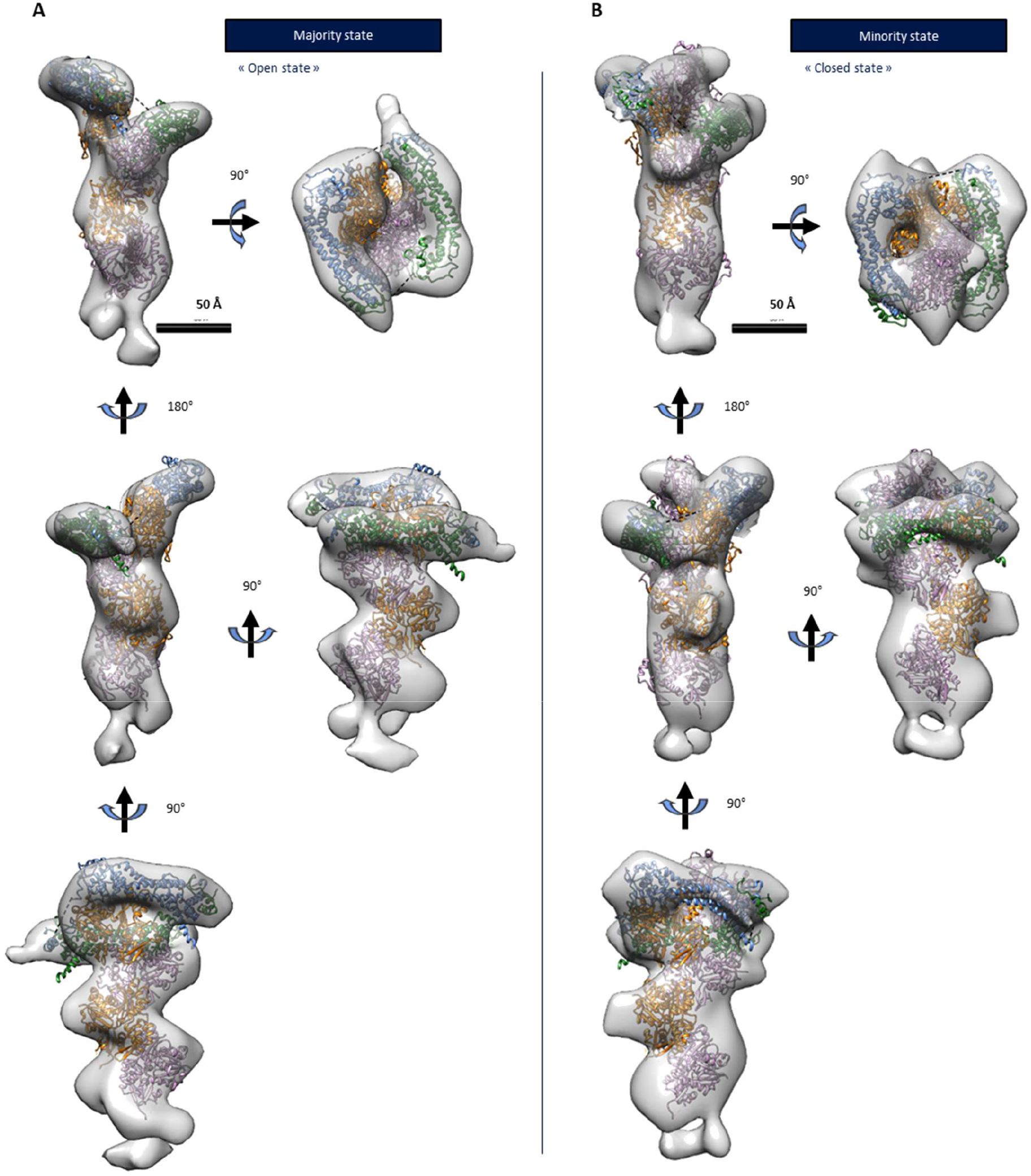
3D structures of formin-bound barbed ends. RELION 3D reconstructions of the majority configuration (**A**, 2048 particles, 29 Å resolution) and the minority configuration (**B**, 402 particles, 32 Å resolution) of the formin-bound barbed ends (grey) in which an atomic structure of the open (**A**) or the closed (**B**) states described by the “stair-stepping” model are fitted (PDB 1Y64/5OOE). *Green/blue*: FH2 domains. *Orange/pink* : actin subunits.

**Figure 3.**
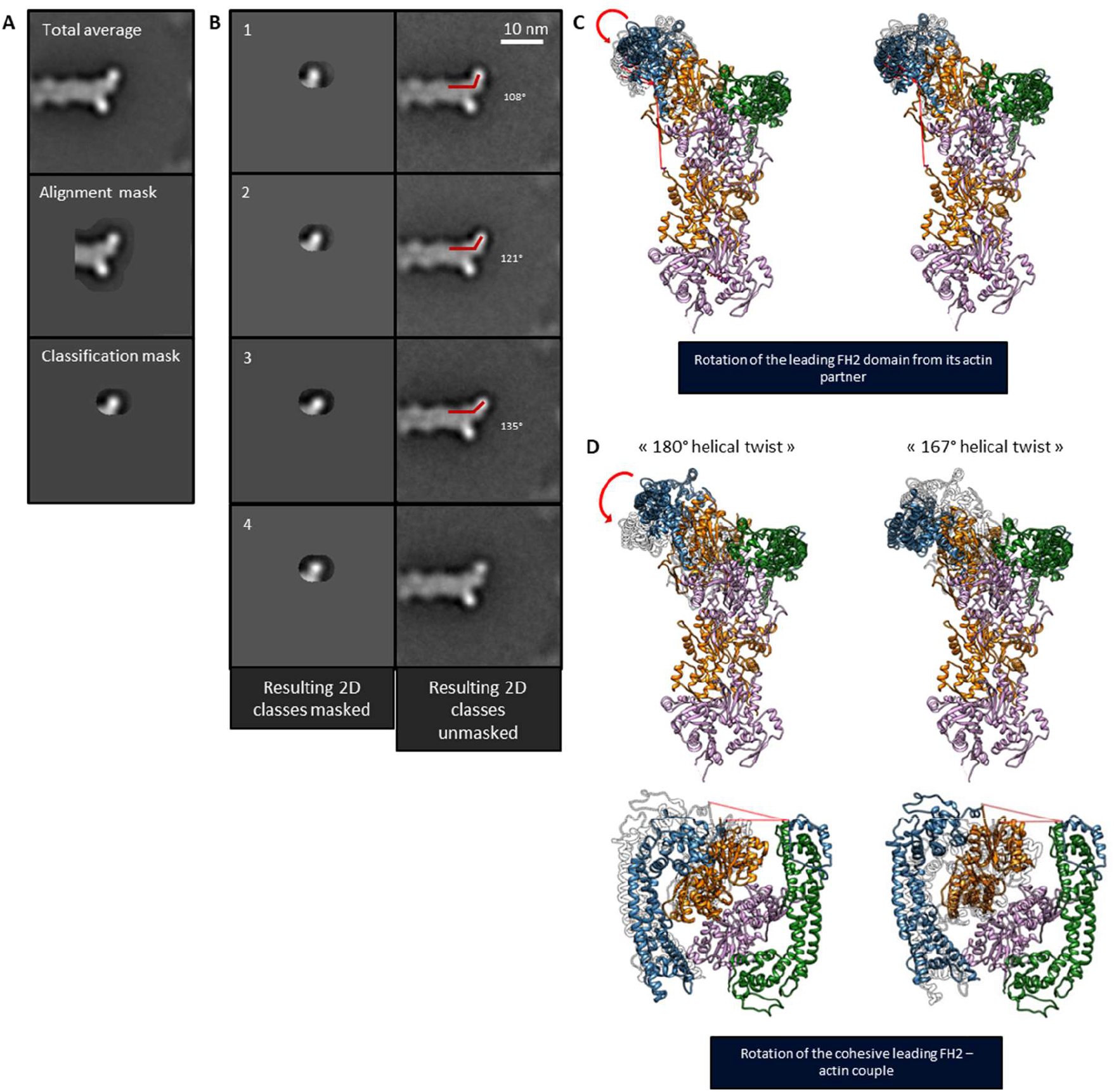
Structural variations in the open state revealed by focused 2D analysis with two hypothetical corresponding 3D models. **A**. *Top*: Total average of formin bound barbed-ends in the open state. *Middle*: Total average displayed through the mask used for particle alignment. *Bottom* - Total average displayed through the focused mask used for particles classification. **B**. Left: classes generated after focused classification and displayed through the focused mask used. Right: classes generated after focused classification and observed without masking. **C**. Atomic models of a barbed end capped with a formin in the open state derived from the previously determined arrangement (see Fig. 2.A) and presented after an independent rotation of the leading FH2 domain in the trigonometric (left) or anti-trigonometric (right) direction. For each two positions shown, the opposite FH2 position is displayed in transparency. **D**. Atomic model of a barbed end capped with a formin in the open state (left) and presented in the previously determined arrangement (see Fig. 2.A) or after reorientation of the first actin subunit (orange) (top-right). Top: side view. Bottom: top view. Green/blue: FH2 domains. Orange/pink: actin subunits. C-D. The red bent arrows describe the rotation movement behaved by the leading FH2 domains while straight red lines materialized angles formed by the filament axis and the FH2 orientation in different configurations.

**Figure 4.**
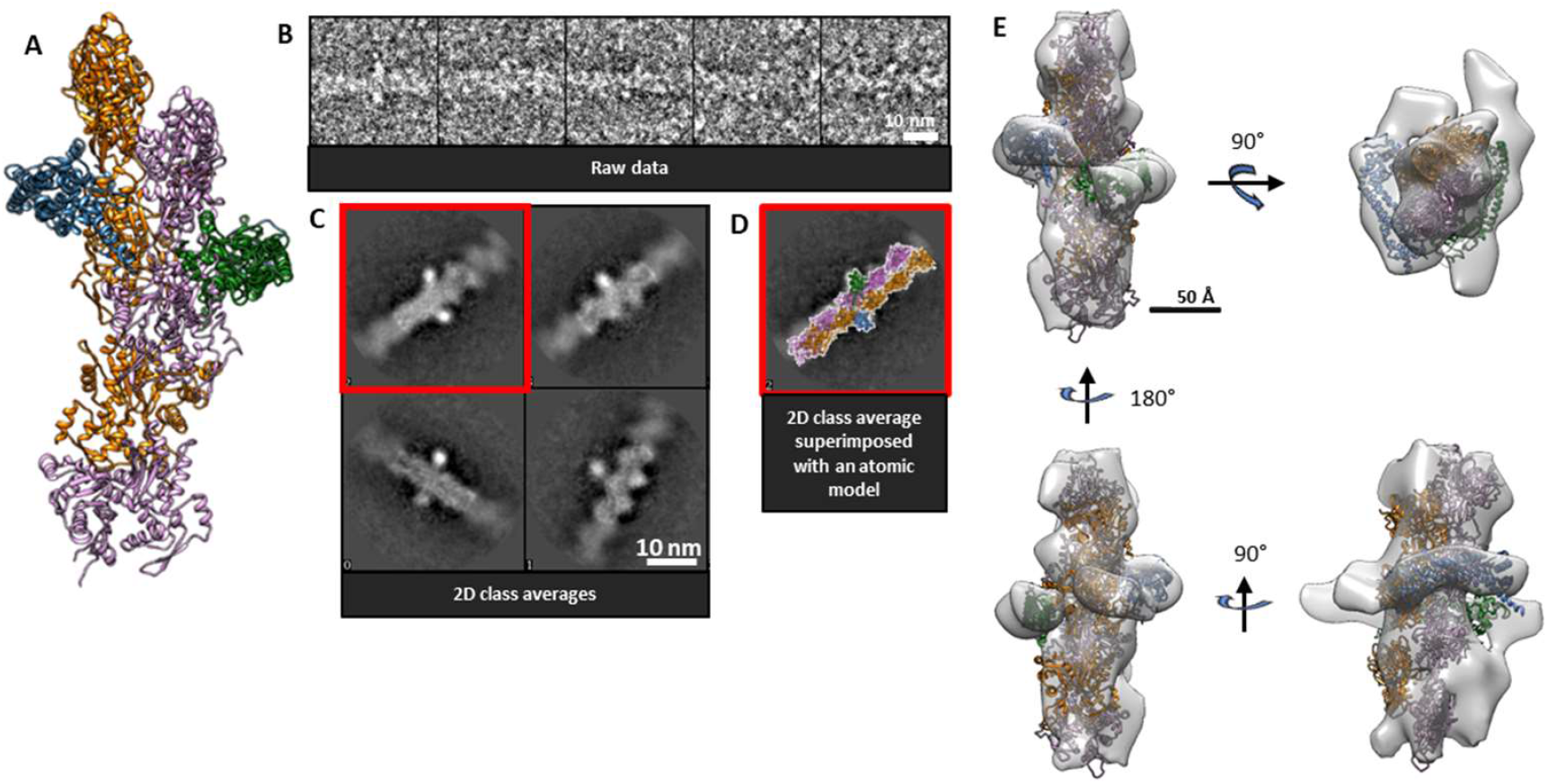
Formin interaction mode with actin filament by encircling the helical body. **A**. Atomic model of an FH2 dimer encircling an actin filament ‘slightly’ away from the barbed end location. This atomic model was constructed according to the open state of the stair stepping model where the leading FH2 domain has been moved away from the filament axis (See Fig 3.) and two actin subunits have been added at the barbed end. **B**. Raw images of particles identified as FH2 domains encircling the actin filament. **C**. RELION 2D classes generated from particles picked along actin filaments previously sonicated and incubated with formins. **D**. Class average (from C. red window) matched with the 2D pattern of an FH2 dimer encircling the actin filament body. **E**. 3D reconstruction of a formin encircling the actin filament body (717 particles) in which atomic structures are fitted (PDB 1Y64/5OOE). *Green/blue*: FH2 domains. *Orange/pink*: actin subunits.

## Results

To observe formins at actin filament barbed ends, we first optimized the density of short actin filaments and thus the number of actin ends adsorbed onto an electron microscopy grid, as described in Supplementary Figure 1 and in the Methods section. Sonication was carried out on 1 μM rabbit alpha-skeletal preformed actin filaments in F-Buffer at pH 7.8 with 50 mM KCl, in order to shorten them before exposing them to formin mDia1 (see Methods). Sonication has been reported to induce the depolymerization of actin filaments (F-actin), in addition to their severing^22^. In order to quantify the amount of actin monomers (G-actin) generated by our sonication step, we performed measurements using pyrene-labeled actin (Supplementary Figure 2) with samples prepared the same way as for EM. They allow us to determine that, when we fixated our samples for EM observation, there was 250-350 nM G-actin in solution, and thus mDia1-bearing barbed ends were elongating at a rate of 1-2 subunits/second.

### Two-dimensional single particle image processing of actin barbed ends

As compared with SPA (Single Particle Analysis) performed on globular proteins, our objects of interest are circumscribed to the ends of elongated asymmetrical assemblies whose upstream bodies extend beyond the periphery of the extracted boxes. Raw boxed particles and an example of filamentous ends displaying an additional density are presented in Figure 1.A-B. We thus adapted single particle analysis to our specific sample to optimize the analysis of the filament ends. The two-dimensional analysis is schematically summarized in Supplementary Figure 3 and presented in the Methods section. After a first round of the alignment protocol, a 2D classification was performed to select the particles corresponding to actin filaments interacting with formins. A second round of the same alignment protocol was performed again with the dataset extracted from the classes displaying a specific signal at the actin ends (highlighted in green in Figure 1.B). At this stage, an additional 2D classification was carried out to improve the resulting 2D classes.

From the resulting 2D analysis, three distinct groups can be identified. From the first round of classification, most of the generated classes correspond to bare actin filament ends (Figure 1.C). These classes are identical to the classes obtained for an actin control sample without formins (see Supplementary Figure 4). These strictly naked actin filament ends represent about 57 % of the dataset (11,960 out of a total of 20,919 particles). These classes were not considered for the second round of classification. From a second round of classification for filaments potentially displaying formins (Figure 1.C, green boxes), two additional groups (see Figure 1.D-E, highlighted in red and orange) clearly display additional densities with determined shapes at the actin filament ends. The classes highlighted in red show a stereotypical “Y shape” and represent 26% of the total dataset if we consider only averages with sharp and defined details (Figure 1.D, continuous red boxes, 5,575 out of 20,919), to 31% of the total dataset if we include averages displaying blurry features (Figure 1.D, continuous and dashed red boxes, 6,356 out of 20,919). The data from the classes showing blurry features (5%) were not considered for further data processing. The corresponding global average can be superimposed with a 2D projection of an atomic model structure of the open state of the “stair stepping” model (Figure 1.E, red box). Atomic models were built using previously published actin and formin FH2 crystallographic structures (PDB: 5OOE and PDB: 1Y64, respectively). The classes highlighted in orange are in minority representing 8% of the total dataset (1,731 particles out of 20,919) and the corresponding global average can be superimposed in 2D with an atomic model structure of the closed state of the “stair stepping” model (Figure 1.E). The proportion of ends in the open state among all the ends where a formin can be visualized adds up to 76-79%, depending on whether one considers averages with or without blurry features. To go further, the aligned particles belonging to the classes corresponding to either the open or the closed state, and displaying sharp details, were merged to be finally subjected to 3D classifications.

### Open and closed conformations of FH2 mDia1

To characterize the possible different conformations in 3D, we have performed multi-reference 3D classifications and reconstructions merging the two sets of particles displaying additional density at actin filament ends within the 2D analysis (see Figure 1, red and orange boxes, 7,306 particles). As initial references, high resolution structures of a bare actin filament barbed end (PDB: 5OOE) and open and closed states of the “stair-stepping” model (generated using PDB: 1Y64) were selected and filtered to low resolution (50 Å). Three 3D classes were obtained during the first round of 3D classification. Two classes reflect two distinct conformations of formin bound to the barbed end of actin filaments. A third class corresponds to misaligned particles. The sub-dataset corresponding to the first two 3D classes were independently subjected to further 3D classification followed by 3D refinement and post-processing. The resulting 3D reconstructions are shown in Figures 2A and 2B, gathering respectively 3,694 (18%) and 977 particles (5% of the total dataset). They reach a resolution of 26 Å and 27 Å, respectively (Supplementary Figure 9). The first class gathers 79 % of formin-bound filament barbed ends identified by 3D classification while the second class gathers 21 % of these formin-decorated barbed ends.

Within these 3D envelopes, a segment of actin filament extends towards the pointed end as already shown by the 2D classes. At the barbed ends, some additional density can be visualized which displays the shape of an FH2 formin dimer. To confirm and clearly delineate the nature and relative orientation of the proteins within our 3D classes, we have carried out automated rigid docking of high-resolution structures in our envelope. The PDB 1Y64 crystallographic structure^1^ of an actin subunit in complex with a single Bni1p FH2 formin domain was docked into the 3D structures. The “Fit in Map” tool from Chimera^23^ was used to dock a doublet of actin-formin complex. In this high-resolution starting model, the actin subunits bound to FH2 domains were set at a helical twist of 180°. Two additional actin subunits from the PDB 5OOE structure^22^ were docked towards the pointed end, with a canonical helical twist of 167°. Following this “rigid” global docking, a local adjustment was performed by a “sequential fit” performed with Chimera to finely dock the FH2 domains independently from one another. The helical twist of the actin subunits towards the pointed ends and the actin-FH2 (“knob”) contacts are imposed and static in the docking process. The resolution of the envelope obtained following this procedure does not allow one to decipher the helical twist adopted by the two terminal actin subunits at the barbed end. Nevertheless, the first 3D class (Figure 2.A) representative of most of the particles (79 % of formin-bound filament barbed ends) closely matches the open state described by Otomo and colleagues^1^. Indeed, the FH2 domains, highlighted in blue and green, are bound to only two actin subunits and unambiguously show one free and accessible “post” domain that could bind a third actin subunit. The fact that this conformation corresponds to 79% of the identified barbed ends with a bound formin strongly argues in favor of the two state “stair stepping” model (Figure 5, top left-hand corner) as a conformation where the post domain of the leading FH2 is not interacting with any actin subunits does not exist in the “stepping second” model^14^. The second 3D class (Figure 2.B) contains an additional actin subunit protruding from the FH2 dimer at the barbed end. In this conformation, the FH2 domains encircle three actin subunits simultaneously, matching the closed state of the “stair stepping” model.

**Figure 5.**
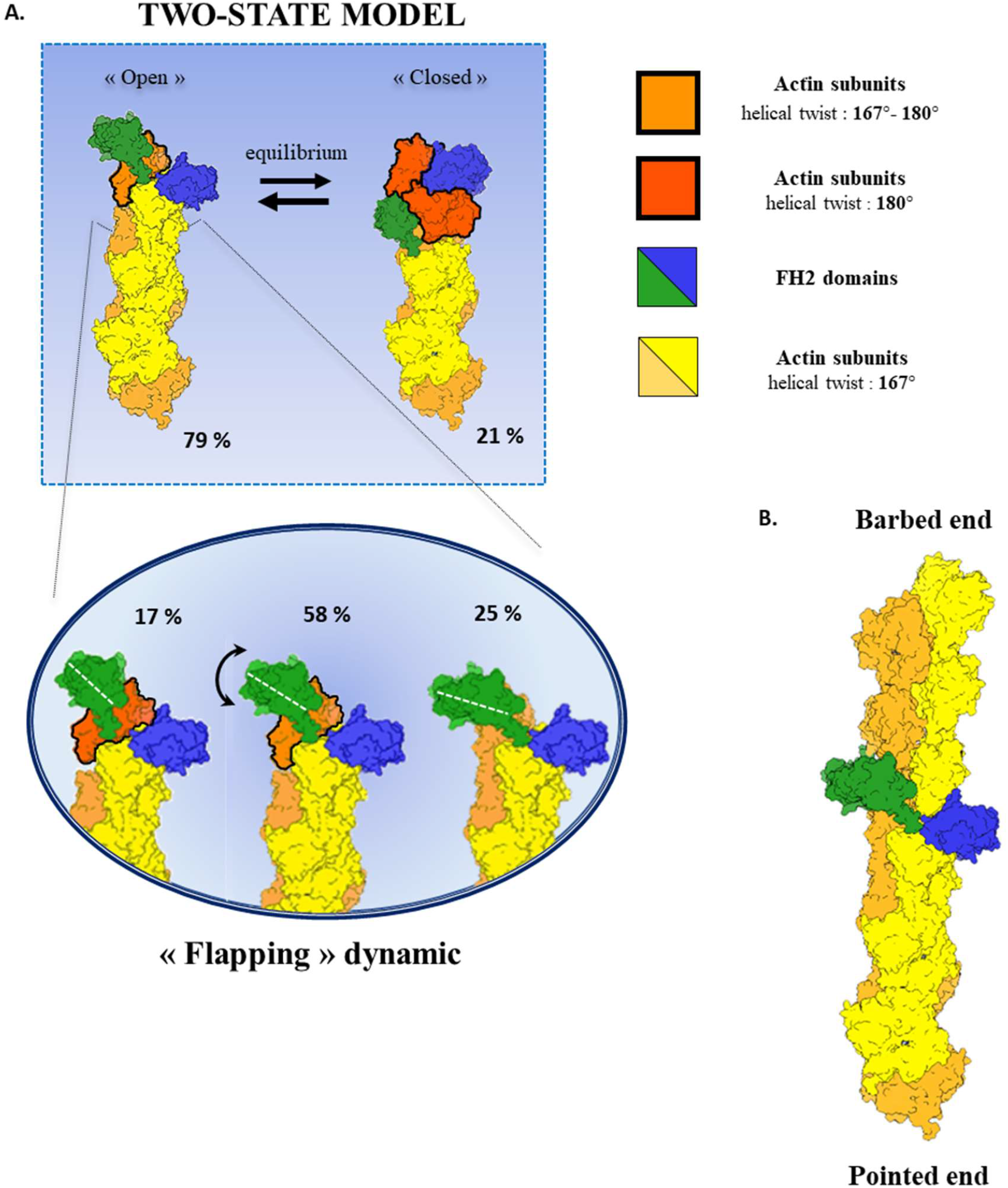
Schematic representation of formin FH2 domains bound to actin. **A. Top:** Schematic representation, based on our 3D reconstructions, of the two-state model proposed by the stair stepping model with percentages indicating the relative distributions of the open and closed states as determined in Otomo et al.^1^. **Bottom:** Schematic representation of the flexibility observed in the open state, with the relative distributions of the main sub-states with percentages indicating the distribution between the main substates. The two strands of the long-pitch double helix are shown in two colors, orange and yellow, for readability. Each of these subunits makes a 167° angle with its nearest short-pitch neighbor. Actin subunits at the barbed end displaying an angle close to 180°, with respect to their short-pitch neighbor, are shown in red. Actin subunits at the barbed end displaying an intermediate angle, between 167° and 180°, with respect to their short-pitch neighbor, are shown in dark orange. The FH2 domains are shown in green or blue. The color code is displayed on the right hand side of the scheme. B. Schematic representation of FH2 domains encircling the core of an actin filament. The color code is identical to A.

To ensure that formins were also visible at the barbed ends of actin filaments in a more physiological condition and were not artifacts from staining and drying by negative stain electron microscopy, we have carried out cryo-EM experiments. To enhance the density of filaments on the cryo-grids, we have used C-flat grids coated with a layer of graphene as described by Palovcak et al.^24^ (see methods section). After sample cryo-fixation, we have collected data using a 200kV FEG “Glacios” microscope. Typical images are displayed in Supplementary Figure 5. From 3,405 images, 12,112 ends were manually picked. Given that the signal-to-noise ratio and the density of ends are lower for cryo-EM images, the final resolution obtained from these data was unfortunately not better than what we obtained with negative stain images. Nonetheless, in 2D classes, additional densities can be pinpointed at the ends of filaments (See Supplementary Figure 5). Due to the limited number of ends observed, we identified primarily one conformation of formin bound to the barbed ends. From 3D reconstruction and classification, one main structure was generated (Supplementary Figure 5C). Actin subunits as well as formin FH2 domains were docked within this structure. The FH2 dimers connect only the two terminal actin subunits of the barbed end with a free and accessible post domain, a recognizable feature of the open conformation.

### Conformational flexibility of the open conformation

After the generation of two-dimensional classes, we noticed that the open state class (see Figure 1, red) displayed some striking variability in the orientation of the FH2 domains with respect to the actin filament. To further analyze this variability, we applied a classification mask centered on the leading FH2 domain (see Figure 3A). Using this mask, the particles found to be in the open conformation (3,804 particles) underwent a classification procedure. The orientation of the leading FH2 domain unambiguously varies as shown in Figure 3B. The amplitude of the angular fluctuation was assessed using ImageJ (see Methods and Supplementary Figure 7). Four classes were obtained displaying three different orientations of the leading FH2 domain (Figure 3B-C). In class 1 (25 % of the particles), the angle between the actin filament long axis and the leading FH2 domain is the lowest (108 degrees). In classes 2 (29 % of the particles) and 4 (29 % of the particles) similar intermediate orientation of the FH2 domains (121 degrees) are found. The details featured in class 4 do not appear as sharp as in class 2, suggesting that within class 4 the conformation slightly varies around the 121° orientation. In class 3 (17% of the particles), the angle between the leading FH2 domain and the actin filament axis is the most obtuse (135 degrees). The amplitude of the fluctuations likely reflects the existence of a continuum of conformations. Indeed, tuning the threshold for classification can generate more classes whose orientation only slightly differs from one another. We chose four classes to make the conformational change clearly visible. In those 2D classes, the actin filament holds the same orientation. The analyzed variability therefore does not result from different views and orientations of the sample. Moreover, the Euler angle distribution assigned during the previously shown 3D reconstruction was analyzed regarding the different 2D classes shown here and their belonging particles. No specific orientation trend could be discerned within the observed 2D classes. Given the small number of particles combined per 2D class and the hypothesized underlying conformational continuum, the generation of 3D structures for these intermediate conformations was not considered.

### FH2 domains encircling the actin filament body

Using single molecule fluorescence microscopy, it was previously shown that mDia1 formins can be displaced from actin filament ends by a capping protein, and formins were then observed to perform 1D diffusion along the actin filament core^25^. Within our images obtained in the absence of capping proteins, we could indeed pinpoint additional densities alongside actin filaments (see Figure 4B). We started our analysis with 2,013 extra densities manually picked and subjected to a 2D classification. The 2D classes are presented in Figure 4C. Additional densities on the side of the filaments are highlighted. The superimposition of high-resolution FH2 and actin structures onto one of the 2D classes (Figure 4D) suggests they are indeed FH2 domains. The particles from the last class (717 particles) were used to generate a 3D structure (Figure 4E). Some distortion, visible on the 3D reconstruction, resulted from a predominance of views and thus of available angles in the 3D reconstruction. Indeed, automated particle picking was unsuccessful and manual picking induced the selection of preferential views.

Nonetheless, we reconstructed a 30 Å resolution structure where the FH2 domains as well as the actin subunits can be pinpointed, and from which an FH2 dimer can be docked encircling an actin filament. The low resolution of the reconstruction is sufficient to show that, alongside actin filaments, the relative arrangement of the FH2 domains and actin is comparable to the ones observed at the actin barbed ends. Our observations thus suggest that FH2 dimers can diffuse along actin filaments by interacting with specific binding sites. We estimated that we could identify on average approximately 1 formin per 4 μm of actin filament (Methods, Supplementary Figure 8).

## Discussion

Using EM, we have, for the first time, directly imaged mDia1 FH2 dimers in interaction with polymerizing actin filaments. The medium-resolution conformations obtained in this study allow us to discriminate between two previously proposed models^1,14^ describing the conformations of formins interacting with growing barbed ends, in favor of the stair-stepping model. Indeed, we clearly observe a conformation where one FH2 dimer interacts with only two actin subunits, and where the leading FH2 domain partly hangs in solution. This conformation is predicted by the stair-stepping model and is completely absent from the stepping-second model.

Nevertheless, our structures have a limited resolution so that finer structural details remain to be uncovered. For instance, we could not directly determine whether the actin subunits of the barbed ends are rather organized with a canonical 167° or a nonstandard 180° helical twist as suggested earlier^13^. In Otomo et al^1^, the 180° non canonical helical twist observed in the crystal might result from the presence of FH2 bound in a “daisy-chain” manner to all of the actin subunits along the filament. It would also be sensible, in a future investigation, to determine actin helicity with formins bound alongside the core of actin filaments.

Compared to recent publications describing pointed ends or capped barbed ends at higher resolution^26^, the dynamic nature of formins at the barbed end makes the production of a high density of short filaments very challenging. In its primary function, capping protein CapZ restricts actin polymerization and thereby permits the production of a high density of short-capped filaments that can be isolated through size exclusion chromatography, and subsequently further concentrated^26^. In our case, to obtain a high density of short filaments, in addition to sonication, the use of additional actin-binding proteins to either sequester monomeric actin in solution (see supplementary Figure 2.C) or to enhance actin nucleation did not lead to any significant improvement of the quality of the data. Our attempts at using additional proteins to trap G-actin (Gc-globulin, Fig S2C) or enhance nucleation (spectrin-actin seeds) did not provide a higher density of formin-bearing barbeds end per field of view. Sonication, our most efficient strategy, was not efficient enough to produce a higher barbed end density that would lead to higher-resolution structures.

A widely-used characteristic of formins is their so-called ‘gating factor’. It is defined as the ratio between the elongation rate of formin-bearing and the elongation rate of bare barbed ends, in the absence of profilin. Formin mDia1 has a notably high gating factor of 0.9. The gating factor is often assumed to represent the fraction of time that the formin spends in the open state. However, this assumption relies on the hypothesis that the monomer on-rate is the same for a bare barbed end than for a formin-bearing barbed end in the open state, which appears very unlikely. Kinetic measurements thus provide limited insights into the relative time spent by a formin in open and closed state.

Alternatively, in an earlier study^15^, by measuring the elongation rate of mDia1-bearing barbed ends as a function of mechanical tension, we have estimated that the FH2 dimer was in the open state 56% of the time, in the absence of tension. This estimation is lower than the outcome of the structural assay presented here. This estimation relies on the consideration that the applied tension skews the state occupancy in favor of the open state, and the experimental data was fitted based on the hypothesis that tension is applied equally to both FH2 hemi-dimers. With an alternative analysis, considering that the force is applied to only one of the two FH2 hemi-dimers (see Supplementary text) we would obtain an even lower estimation (12%) of the open-state occupancy rate in the absence of tension. In these former experiments, filaments were elongated in the presence of profilin-actin, which binds to polyproline tracks of the FH1 domains to deliver actin to the barbed end. The simultaneous interaction of profilin-actin with the FH1 domain, and with the FH2-bound barbed end, transiently forms a ‘ring complex^17,27^. One cannot exclude that the formation of this ring complex might decrease the open state occupancy rate, compared to the absence of profilin.

Our direct assessment of the open state occupancy rate thus provides important information on the molecular nature of the formin-barbed end conformations which could not be directly inferred from kinetic measurements, with or without mechanical tension, so far. Considering a gating factor of 0.9 and considering that formin mDia1 spends 79% of the time in the open state, we can compute that the on-rate for monomers would be slightly higher (14% higher) for an mDia1-bearing barbed end in the open state, than for a bare barbed end. The available actin-binding interface on the leading FH2 domain likely provides a first docking intermediate for actin monomers that would help their orientation relative to the barbed end, resulting in a higher on-rate.

We also reveal that, in the open state, the orientation of the leading FH2 relative to the long axis of the filament fluctuates (Figure 3). Several conformations can be distinguished, suggesting a continuum of conformations. These conformations are likely to have different on-rate constants kon for the addition of actin subunits at the barbed end. This could partially explain why previous measurements, based on assembly rates, estimated a lower occupancy rate for the open state^15^.

We propose a “flapping” model where the angle between an FH2 domain and actin varies from 108 to 135 degrees (Figure 5). We designed extrapolated 3D models from existing high-resolution crystal structures that would exhibit, in 2D projections, the observed angular “flapping” fluctuations (Figure 3). In a first attempted model, one can examine whether the rotation of an FH2 domain independently from the filament barbed end structure would be possible. As shown in Figure 3.C, the rotation is not constrained by any steric clash and would be allowed by the flexibility of the FH2 “linker” domain. However, upon rotation, the FH2 “knob” domain might not interact properly anymore with the terminal actin subunit. Hence, a second more favorable model is proposed (Figure 3.D) where the terminal actin subunit would rotate with a slight bow and thus drag an FH2 interacting through its “knob” domain. In such a configuration (Figure 3.D), the contacts between the FH2 domain and actin subunits would thus be preserved and their relative orientation and spacing imposed. Following this rearrangement, the actin helical twist would vary in a range included between the two observed values of 180° (structure 1Y64) and 167° (structure 5OOE) similarly to the “stepping second” mechanism^14^ and to what was consequently assessed by Molecular Dynamic simulations^16^. However, our data exclude the “stepping second” model to describe FH2 dimer translocation in actin polymerization. The terminal actin subunits at the barbed end would thus show a configuration closer to the helical twist angle found within canonical actin filaments and the previously characterized formin-actin contacts in 1Y64 would be preserved. Nonetheless, while this angular switch cannot be directly observed within our medium-resolution structures, the proposed mechanism is a sensible hypothesis that would reflect the “flapping” FH2 domain motion towards its most open conformation. Indeed, the two actin filament arrangements considered as the extreme configurations of such a motion (PDB structures 1Y64 and 5OOE) differed not only by their helical twist value but also by the actin subunits’ orientations relative to the filament axis^1^ resulting in actin subunit rotating outwards within the most open conformation. In this extreme open conformation, after the transition from an actin helical twist of 180° to 167°, an actin monomer (polymerization) or an actin oligomer (annealing) could be added at the barbed end in the canonical actin filament conformation (Supplementary Figure 10). Indeed, actin polymerization occurs following sonication in our experimental conditions (Supplementary Figure 2), and annealing cannot be excluded.

From this state, and without any actin twist back to 180° occurring before the addition of additional actin subunits, the FH2 dimer would then be “lagging” behind the elongating barbed end, without being synchronously translocated with it (Supplementary Figure 10). Lagging formins could then start diffusing along the actin core, in both directions, and such a scenario would result in formins encircling the core of actin filament away from the barbed ends. To our knowledge, such a situation has never been observed, in the absence of capping proteins. Nonetheless, the observations from Bombardier et al.^25^ suggest that formins can be displaced from the barbed end to the core of actin filaments, and our results indicate that this could happen in the absence of capping protein. This displacement can perhaps be provoked by the binding of a second formin dimer to the barbed end. Other explanations for our observation of formins along the filament include the direct binding of formins to the filament sides, which would require the formin dimer to transiently open up.

It was previously observed that the formin dissociation rate increases with the actin filament elongation rate, in the presence of increasing actin concentrations^14,17^. This observation could be partially explained by the ‘lagging’ formin mechanism we propose, where the lagging probability would actually increase with actin concentration. In vivo though, the formin elongation of filament barbed ends are expected to occur primarily from profilin-actin complexes delivered to filament barbed ends mediated by FH1 domains. This mode of elongation imposes frequent interactions between FH1 and the barbed end, which would likely reduce the probability of formins to lag behind the barbed end. In addition, active formins are thought to be anchored to membranes^28^, which should restrict further their ability to diffuse along the core of actin filaments.

In conclusion, our EM observations provide a direct visualization of the different conformational states of mDia1 formins interacting with actin filaments, and refine the processive “stair stepping” model proposed by Otomo and colleagues^1^, by showing a continuum of conformations for the open state. We report here that formins can unexpectedly be found within the core of actin filaments, in the absence of capping proteins that would displace them from the barbed end. We expect that future studies will reveal finer details of the terminal actin subunits arrangement of formin-bearing barbed ends.

## Methods

### Protein purification and storage

Actin was purified from rabbit muscle as detailed previously^29,30^ and stored up to 4 weeks in G-buffer (5 mM TRIS pH 7.8, 0.2 mM ATP, 0.1 mM CaCl2, 1 mM DTT and 0.01% NaN3), on ice.

Recombinant mouse mDia1(SNAP-FH1FH2-DAD-6xHis) formin (uniprot O08808, seq. 552-1255 aa) was expressed in *E. coli* and purified through immobilized metal affinity chromatography (HisTrap) followed by steric exclusion chromatography (HiLoad 16/60 Superdex 200) as developed in Jégou et al.^18^. Fromins were snap frozen and stored at -80°C in 50 mM Hepes buffer pH 7.8, 200 mM KCl, 10% glycerol and 1 mM DTT.

### Pyrene assays

5 μM actin was polymerized at room temperature for 1h in F-Buffer (5 mM Tris-HCl pH 7.8, 1 mM MgCl2, 0.2 mM EGTA, 0.2 mM ATP, 1 mM DTT, 50 mM KCl)to reach steady state. A total volume of 500 μL containing the polymerized F-actin diluted to 1 µM was sonicated in F-buffer. The sonication was operated with a sonicator Vibra-Cell 75041 (20 kHz, 750 Watts, 20% power) equipped with a 3 mm diameter probe. 10 pulsations of 0.1 s each separated by 3.9 s or 4.9 s of rest periods were used for the sonication. 200 to 300 μL of the sonicated solution were then inserted into quartz cuvettes and analyzed for fluorescence signal for at least 20 minutes using a spectrofluorimeter (Xenius, Safas). The average time from the end of the sonication process to the pyrene fluorescence measurement was 30 seconds, in average. As a control, the fluorescence of G-actin and 1 µM F-actin (without sonication) were also assayed.

### Sample preparation for electron microscopy

G-actin stock solution was ultracentrifuged for 45 minutes at 80, 000 g before use to remove any aggregates. 5 μM actin was polymerized at room temperature for 1h in F-Buffer (5 mM Tris-HCl pH 7.8, 1 mM MgCl2, 0.2 mM EGTA, 0.2 mM ATP, 1 mM DTT, 50 mM KCl). To generate a high density of short actin filaments, 500 μL of polymerized 1 µM F-actin was sonicated in F-buffer. The sonication was operated with a sonicator Vibra-Cell 75041 (20 kHz, 750 Watts, 20% power) equipped with a 3 mm diameter probe. 10 pulsations of 0.1 s each separated by 3.9 s of rest periods were used for the sonication. Formin mDia1(SNAP-FH1FH2-DAD-His) was added in the F-actin sonicated mixture directly after the last pulsation, through a Hamilton syringe equipped with a 0.13 mm internal diameter needle. A formin concentration of 100 nM was used for negative stain electron microscopy experiments, and a formin concentration of 125 nM was used for cryo-electron microscopy experiments.20 seconds after formin addition, 4 μL of sonicated F-actin/Formin mixture were collected with a Hamilton syringe. The collected volume was directly applied to an electron microscopy grid.

For negative stain electron microscopy experiments, the solution was adsorbed 30 s on a freshly glow-discharged 300 mesh carbon-coated copper grid. Excess volume was blotted off with a filter paper Whatman n°1. 4 μl of uranyl formiate 1% were transiently added before being blotted off on the grid. A volume of 4 μL of rinsing uranyl formiate 1% was deposited on the grid for 30 seconds before drying it with a paper filter Whatman n°1.

As shown in Supplementary Figure 1.B, short filaments of a few hundreds of nanometers are generated. On average, 13 ends are obtained per image of about 1 µm² (average over 1632 images). In parallel, we have assessed whether generating a density of filaments by using spectrin-actin seeds would sufficiently enhance the density of short filaments at short times. However, the resulting density of short actin filaments remains low and the presence of spectrin-actin seeds, that are not generating filaments, induce an additional background signal which was deleterious to SPA.

For cryo-electron microscopy experiments, C-Flat R 2/2 grids were covered by graphene oxide sheets by depositing 0.2 mg/ml aqueous graphene oxide drops. The grids were used after an overnight drying step as already described^24^. The graphene oxide solution concentration was estimated by spectrophotometer measurements.

4 μL of sonicated actin-F/Formin mixture were incubated for 30 s on these freshly glow-discharged grids. The excess volume was blotted off with a filter paper Whatman n°1 for 3 s before the grids were plunge frozen in liquid ethane for vitrification. These operations were carried out using an automated plunge freezing apparatus (Leica EM GP) operated at 80% humidity.

### Electron microscopy data collection

Negative staining electron microscopy data was collected with a FEI tecnai G2 transmission electron microscope equipped with a LaB6 emission filament operating at a 200 kV acceleration voltage. Images were captured on a TVIPS F416 CMOS camera at a 50,000 magnification and 1.5–2.5 μm underfocus. The corresponding pixel size was 2.13 Å per pixel. A total of 1632 images were acquired.

Cryo-electron microscopy data was collected using a Glacios 200kV (Thermo fisher) transmission electron microscope equipped with a FEG operating at a 200 kV acceleration voltage. Images were captured using a Falcon 3 direct detection camera at ×60,000 magnification and 1–3 μm underfocus. The corresponding pixel size was 2.5 Å per pixel. Each image acquisition was performed in dose-fractionated mode with 60 frames over 2,2 s exposition time for a total dose of 60 e^-^/Å^2^ (1 e^-^/Å^2^ per frame). A total of 4223 movies was acquired.

### Estimation of the average length of actin filament observed per image and density of formins along actin filaments

The actin filament length density observed per image was estimated to extrapolate the actin filament length imaged within the data. The actin filament length in our images is then compared with the density of formins identified along the actin filament bodies to assess the density of formin per actin filament length. The actin filament length was measured manually in 10 images and compared with the measurements resulting from a semi-automated approach on the same images (Supplementary 8). An error of about 5% was estimated between the two approaches. The semi-automated approach was applied to 32 images with a mean length of actin filament of about 4 μm per image. This measurement extrapolated to all the images used for the picking of formins along actin filaments (632 images) leads to a total length of actin filament imaged of about 2566 μm. This length compared to the 717 particles clearly identified as formins encircling the actin filament body leads to an estimation of about 0.3 formin per micrometer of actin filament.

### 2D data processing

From the 1632 micrographs obtained by negative staining electron microscopy, 20 919 actin filament ends were hand-picked using EMAN2^31^ software and extracted with a square box size of 180 pixels^2^ (2.13 Å/pixel). From 632 of these micrographs, 2013 formin-like patterns were hand-picked along the actin filament cores.

A protocol was adapted to the specificity of our data. Dedicated scripts were generated within the SPIDER^32,33^ software. In the first round of alignment, all particles were normalized using a circular mask that roughly eclipses the signal associated with the proteins. In the second SPIDER alignment step, these normalized particles were rotationally aligned and a new normalization step was performed using a rectangular mask that more accurately eclipses the signal of the observed filaments. In a third SPIDER alignment step, a principal component analysis was performed on the rotationally aligned particles. The purpose of this step was to generate reference averages associated with a high signal-to-noise ratio that sample the various global patterns of the extracted particles. The reference averages thus generated were aligned with each other and used for a multi-reference alignment of the particles corresponding to the fourth SPIDER alignment step. The main purpose of this fourth step was to optimize the orientation of each particle. In this operation, a low-pass filter was applied to the particles and the reference averages to prioritize the global orientation of each particle. At this stage, a small adjustment range was allowed for the translational alignment. In a fifth SPIDER alignment step, a new principal component analysis was performed through a mask focused on the tip of the extracted filament ends or focused on the central part of the actin core filament containing a potential formin dimer. The purpose of this step was to generate reference averages representatives of the structural variability and the variability in the positions of the tips. Once again, the reference averages generated were aligned with each other and used for a multi-reference alignment of the particles corresponding to a sixth SPIDER alignment step. The purpose of this last multi-reference alignment was to fine-tune the respective positions of the ends in the first case or of the formin-like pattern along the actin filament core in the second case.

Regarding the actin filament ends, the binding of FH2 domains to the actin filament tips generate asymmetrical 2D patterns with two possible configurations when the FH2 domains are observed through their main axis. The first FH2 domain can appear on either side of the observed actin filament tip with the second FH2 domain on the corresponding opposite side. Both images of a capped filament observed from one side or the opposite side can match with these two configurations. While these two images contain the same 2D structural information, their opposing asymmetric patterns require the application of a symmetry operation along the filament axis to be correctly aligned.

In line with this observation, for each of the particles considered, the alignment score of its mirror particle (symmetry applied along the filament axis) with the reference was also evaluated. In this step, either the raw particle or its mirror particle was kept by selecting the one with the highest alignment score. On this occasion, all the references have been manually oriented in a given direction. This step allows the use of a more focused asymmetric mask around the reference averages and brings together a larger number of particles in the same classes, thus improving the associated signal-to-noise ratios. Given that the number of formin-capped ends that could be identified in the data set will be a minority, this step appears to be important to strengthen their signal.

In the seventh and last SPIDER analysis step, a principal components analysis was performed to roughly distinguish between formin-bound and bare actin filaments. The more homogeneous datasets considered to be formin-bound were then subjected to a second round of 2D SPIDER analysis to benefit this time from the preponderant signal of the formins in the new particle sub-selection.

These aligned data sets were subjected to classical 3D analysis under RELION^34^ v3 in order to benefit from the Bayesian statistical framework and thus retain only the most consistent particles. Finally, 3694 formin-bound filament ends corresponding to the « open state », 977 formin-bound filament ends corresponding to the « closed state » and 717 formin-bound filament cores were determined by RELION 3D classifications.

The 2D classification focused on the first FH2 domain orientation variability among the actin filament ends bound by formin in the « open state » was performed with SPIDER. From 3804 particles, 4 classes were generated with respectively 956 particles (25%), 1092 particles (29%), 651 particles (17%) and 1104 particles (29%). Angle measurements associated with the first FH2 domain orientation in each of the four classes was carried out using ImageJ. Before each measurement, the class average observed through his focused mask was thresholded to distinguish background versus protein density. The edge delimiting background and protein density was calculated with the “Find Edges” ImageJ function. This delimitation was refined with “Skeletonize” ImageJ function. After that, the angle formed by the main axis of the actin filament and the relative orientation of the first FH2 domain was measured.

Movies acquired by cryo-electron microscopy have been processed by MotionCor2 program to correct electron beam-induced sample motion. Contrast Transferred Functions (CTF) from motion corrected micrographs were estimated using the program CTFFIND4. Out of 4233 corrected micrographs, 3405 were retained after eliminating images showing too much drift (broken membrane near the imaged region), aggregates of graphene oxide sheets or proteins, poor confidence of CTF fitting estimation. A total of 12 112 actin filament ends were hand-picked within EMAN2 software and extracted with a square box size of 154 pixels^2^ (2.5 Å/pixel). From this data set and using RELION 2D classification, at least 1548 particles could be identified as actin filament ends bound by a formin dimer.

### 3D Data processing

Structural models of a formin-bound barbed end in his « open state » or his « closed state » and structural model of a formin encircling the actin filament core were generated within UCSF Chimera based on FH2-actin atomic structure (PDB: 1Y64) and actin filament atomic structure (PDB : 5OOE). These low-passed-filtered (50 Å) structural models were used as multireference models for RELION 3D classifications performed on the data sets previously determined by 2D analysis. For each data subset identified after RELION 3D classification, a crude cylinder displaying the adapted dimensions could be used as reference to generate a coherent 3D reconstruction.

After RELION 3D classifications, 3694 particles corresponding to formin-bound filament ends in their « open state » were selected and subjected to a 3D refinement leading to a final 3D enveloppe at a resolution of 26 Å.

After RELION 3D classification, 977 particles corresponding to formin-bound filament ends in their « closed state » were selected and subjected to a 3D refinement leading to a final 3D envelope at a resolution of 27 Å.

From 2013 particles corresponding to a formin dimer encircling an actin filament core, 717 particles were selected after RELION 2D classification and subjected to a 3D refinement leading to a final 3D envelope at a resolution of 30 Å.

For cryo electron microscopy analysis, 2373 particles were selected through RELION 3D classification and leading to a final 3D envelope at a resolution of 30 Å.

In each 3D envelope generated the corresponding atomic model was fitted using UCSF Chimera function « Fit in map » by allowing sequential fitting of independents domain groups. Spurious noise from electron microscopy densities was hidden with the ‘hide dust’ command in UCSF Chimera to facilitate readability.

## Acknowledgments

We thank Daniel Lévy and Manuela Dezi for fruitful discussions. We are grateful to Agence Nationale de la Recherche (ANR) for funding the contract “CONFORMIN” ANR-16-CE11-0013-01. We thank the Labex Cell(n)Scale (ANR-11-LABX0038) and to Paris Sciences et Lettres (ANR-10-IDEX-0001-02). We thank the Cell and Tissue Imaging (PICT-IBiSA), Institut Curie, member of the French National Research Infrastructure France-BioImaging (ANR10-INBS-04). We acknowledge UtechS Ultrastructural3 BioImaging (UBI), supported by the French National Research Agency (France BioImaging; ANR-10–4 INSB–04; Investments for the Future). We thank Stéphane Tachon from NCF (Nanocore facility Imaging, Institut Pasteur).

## Author contributions

J.M., A.B., G.R.L. and A.J designed the experiments. J.M., AB. and A.DC performed the EM experiments. B.G. purified the proteins. A.J. and G.R.L. carried out the pyrene assays. J.M., A.B, A.J and G.R.L. wrote the manuscript.

