## supplemental material for "Direct observation of the conformational states of formin mDia1 at actin filament barbed ends and along the filament"

### Supplementary Material

**Supplementary Figure 1. Density of actin filament ends in the presence of formins, with or without sonication treatment**.

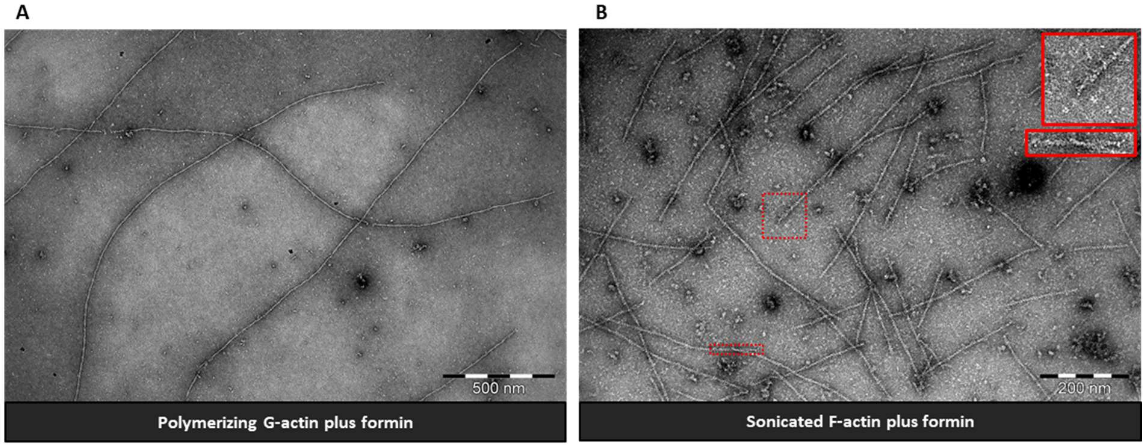


**A**. Actin filaments polymerized in the presence of formins. **B**. Actin filaments grown to equilibrium, fragmented by sonication and incubated with formins before fixation and observation. Inserts (red): actin filament ends exhibiting a typical density contour. Scale bars: A: 500 nm, B: 200 nm.

**Supplementary Figure 2. Pyrene assay after sonication of F-actin**

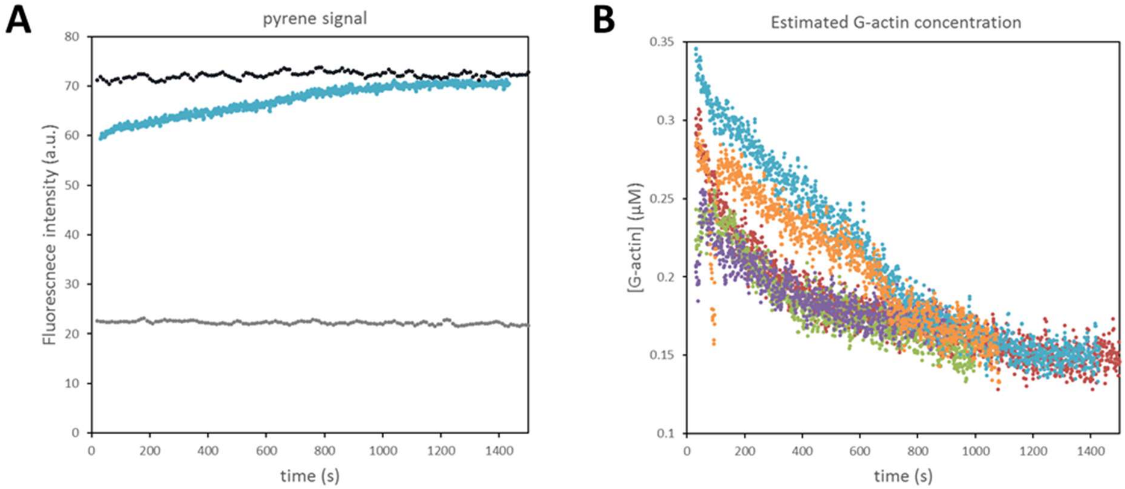


A. Fluorescence signal of pyrene-actin (5% labeled) following sonication, in a sample prepared following the same protocol as the samples for electron microscopy (see Methods) with a final concentration of 1µM actin (blue). The increase of the fluorescence signal indicates that actin is polymerizing. The fluorescence intensity of 1 µM G-actin (gray) and of 1 µM actin at steady-state in polymerizing conditions (black) are shown for comparison.

B. Concentration of G-actin, following sonication, estimated from pyrene-actin fluorescence curves like the one shown in A, considering a steady-state G-actin concentration of 0.15 µM. The curves are 5 independent repeats of the same experiment (the blue curve corresponds to the data shown in A). The rapid variations in the orange and purple curves correspond to transient changes in the fluorescent signal which are likely due to air bubbles or diffusing impurities.

Supplementary Figure 3. Schematic representation of the workflow used to perform 2D image processing of actin filament ends.

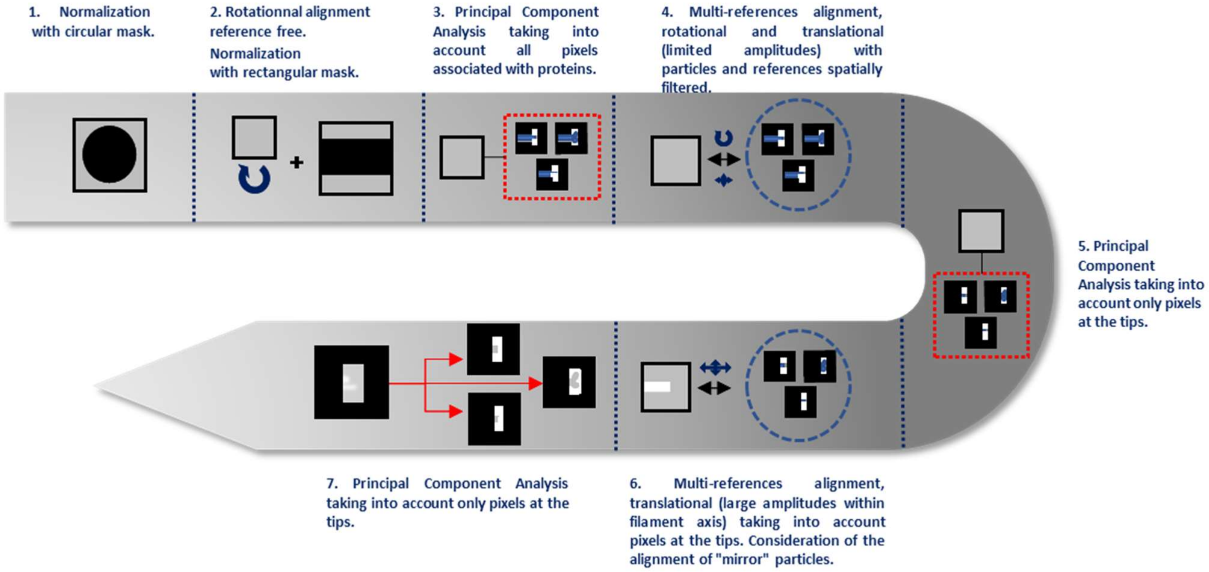

**Supplementary Figure 4. 2D classes obtained from actin filament ends in the absence of any formin. Scale bar 10 nm.**

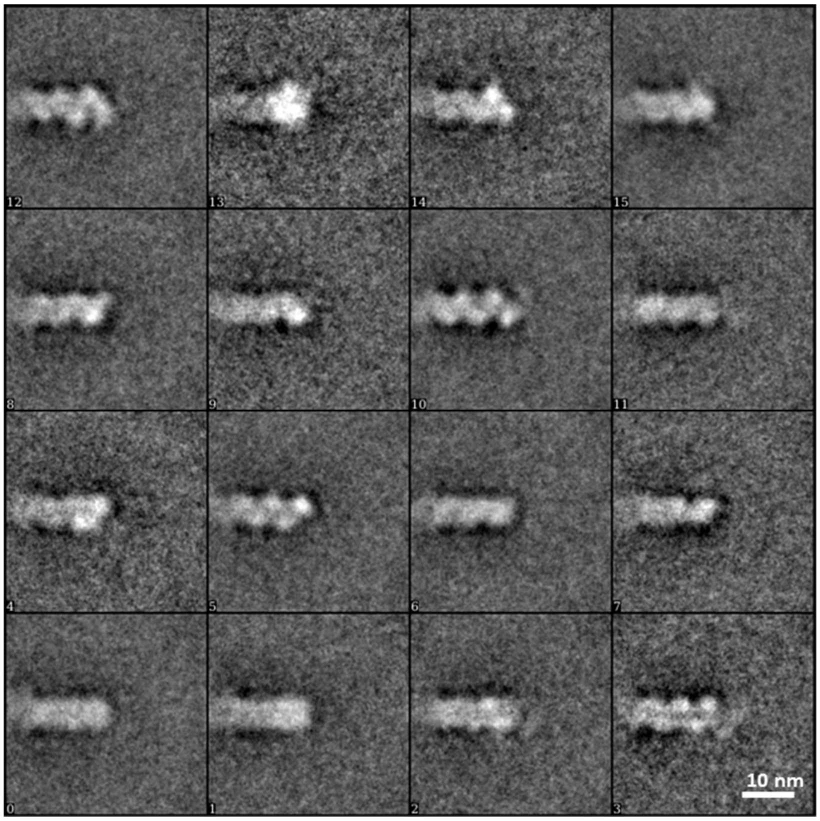

**Supplementary Figure 5. Cryo-EM image of actin filaments observed by cryo-electron microscopy after actin filament sonication followed by incubation with formins.**

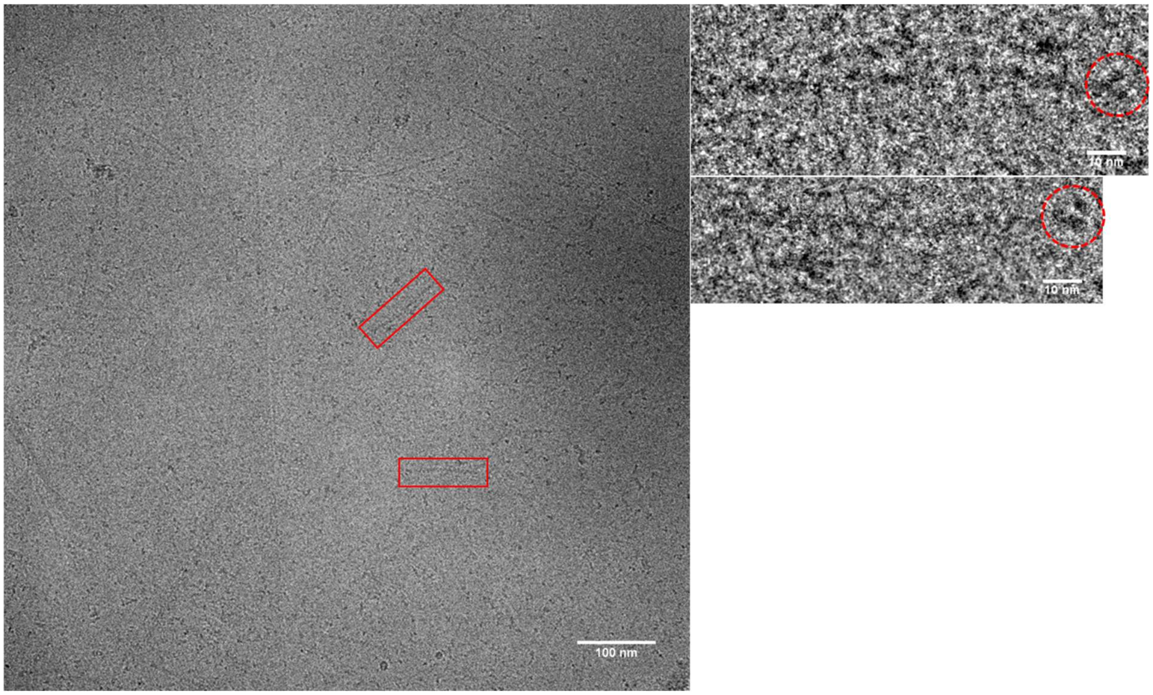


Two actin filaments unambiguously decorated at one end are highlighted with red boxes (Scale bar: 100 nm). Enlargements of these ends are displayed in the inserts where ends densities are point out by dashed red circles (Scale bar: 10 nm).

**Supplementary Figure 6. Cryo-electron microscopy evidence of formin-bound barbed ends.**

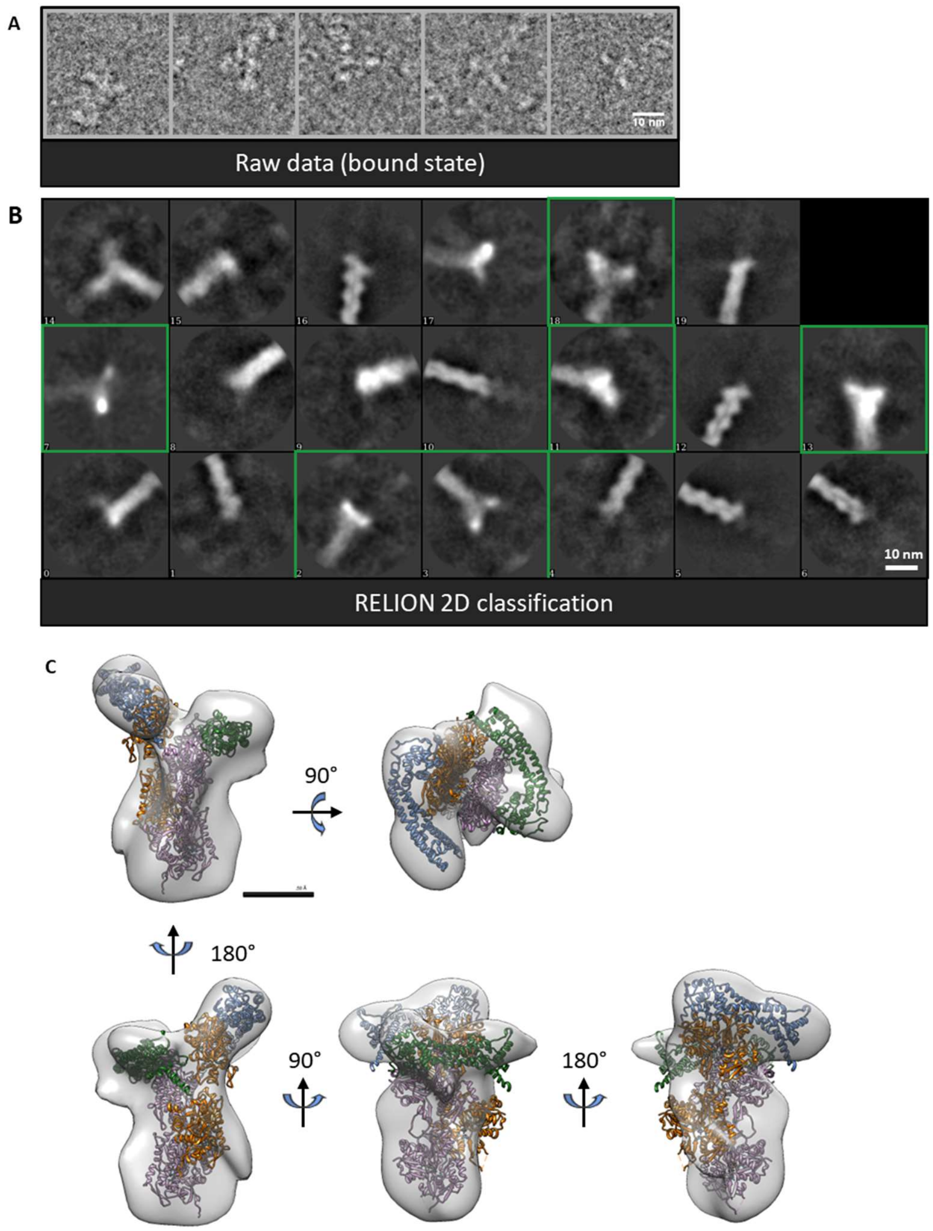


**A.** Raw images corresponding to actin filament ends observed by cryo-electron microscopy after actin filament sonication followed by incubation with formins and showing additional densities. **B.** RELION 2D classes generated from actin filament ends (12 112 particles). *Green windows* : 2D classes of actin filament ends showing additional densities attributable to bound formins. **C**. 3D reconstruction of formin-bound barbed end (2283 particles) in which an atomic structure of the "open" state described by the "stair-stepping" model is fitted (PDB 1Y64/5OOE). *Green/blue*: FH2 domains. *Orange/pink:* actin subunits.

**Supplementary Figure 7. 2D measurement procedure of FH2 domain movement amplitude at the barbed end in the open state.**

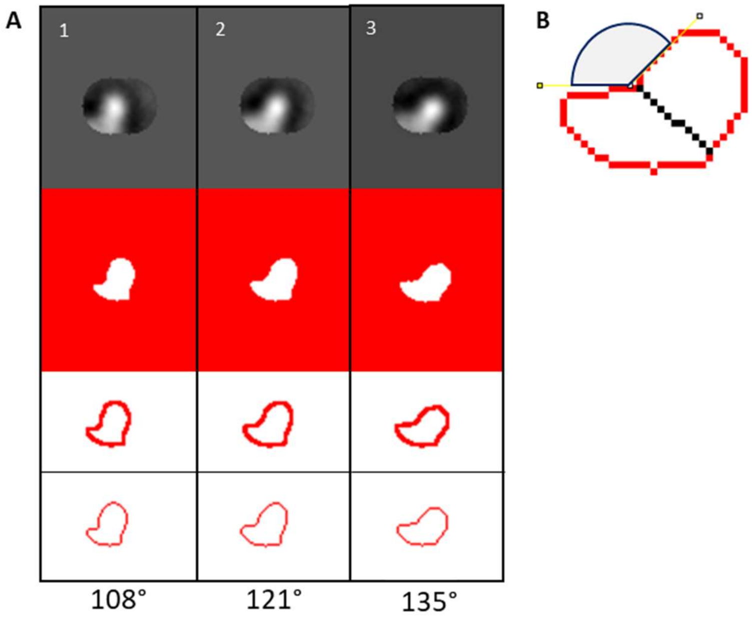


Left, middle, right: class average n°1, n°2, n°3. **A**. Row n°1 : Masked class averages. Row n°2: Binarised class averages. Row n°3 – Filament end border delineation. Row n°4: 1D edge reduction. Row 5 - Angle values measured. **B**. Principle of measuring an angle from the generated 1D border

**Supplementary Figure 8. Methods used to segment actin filaments and determine the length of actin per image as presented in the methods section**

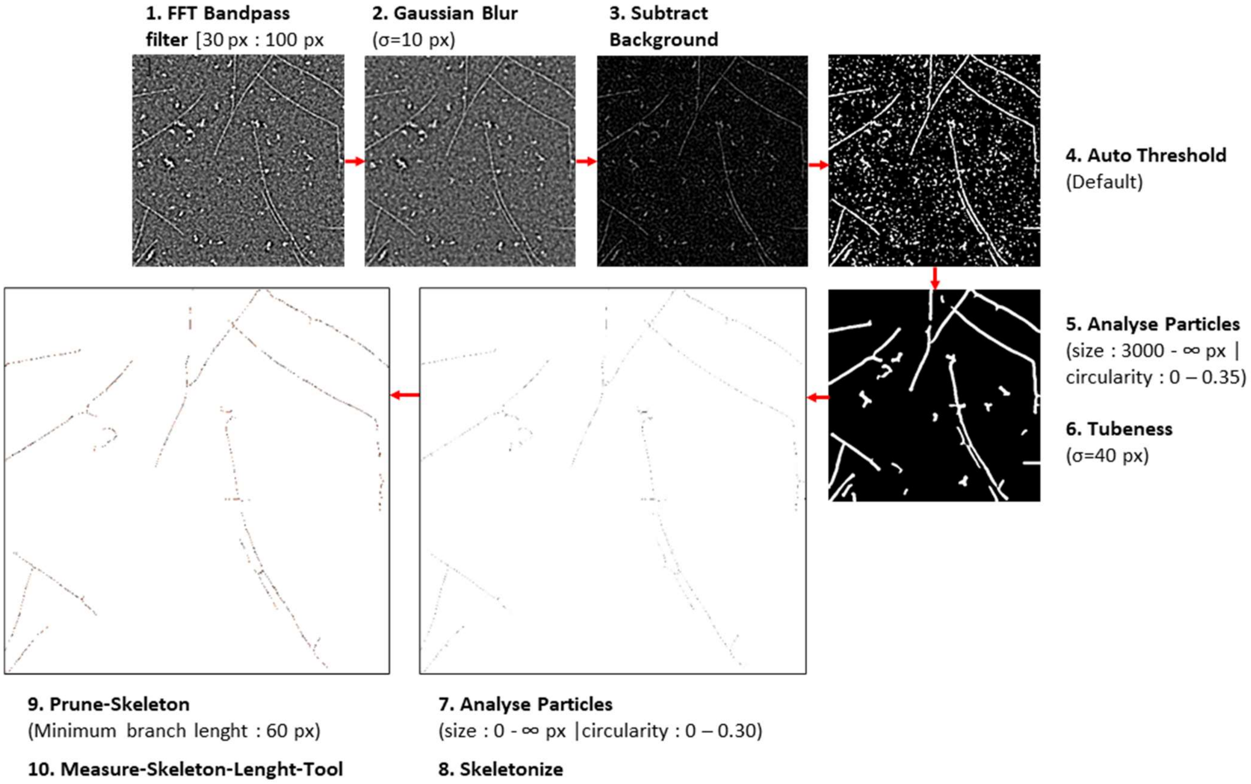

**Supplementary Figure 9.**

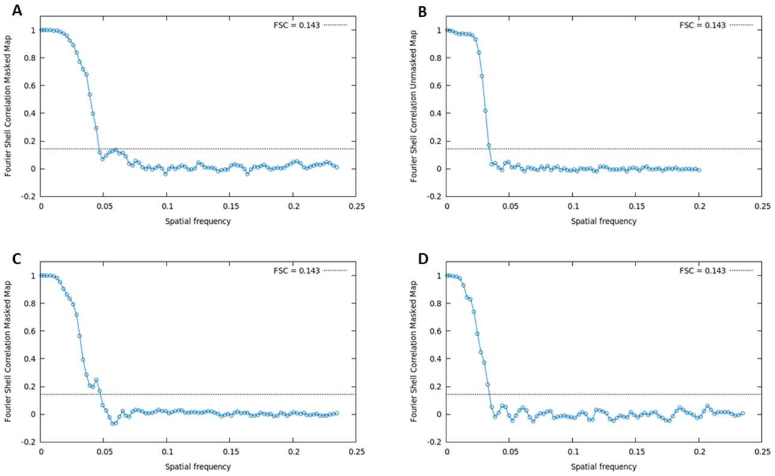


FSC associated to 3D reconstructions of formin-bound barbed ends in the « open » state, observed by negative staining electron microscopy (**A**) or cryo-electron microscopy (**B**). **C**. FSC associated to 3D reconstruction of formin-bound barbed ends in the « closed » state and observed by negative staining electron microscopy. **D**. FSC associated to 3D reconstruction of formin-encircled actin filament observed by negative staining electron microscopy.

**Supplementary Figure 10. Hypothetical scenarios placing the formin in the core of the filament.**

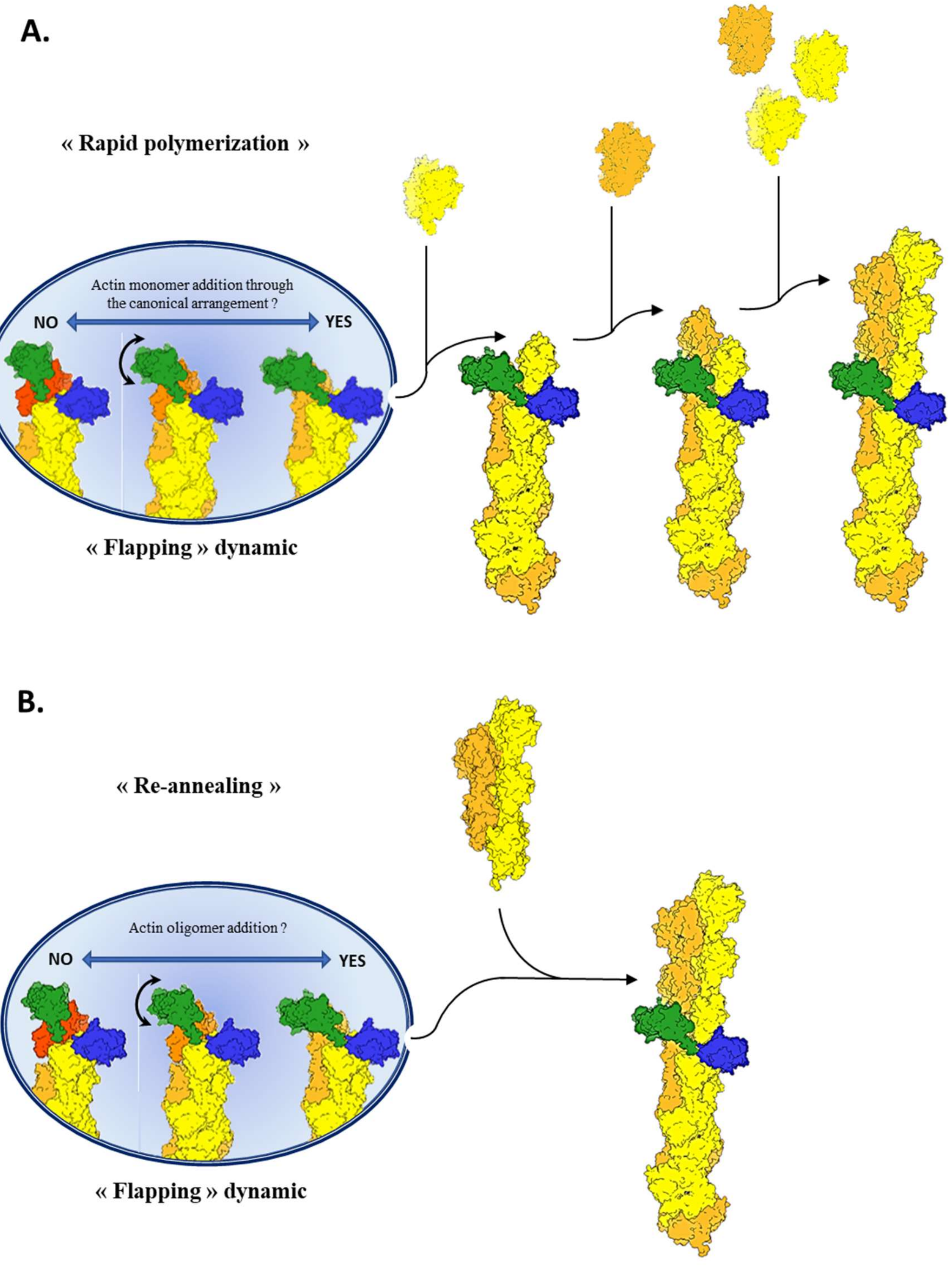


A.Sequential addition of individual actin monomers (orange and yellow) at the barbed ends, leaving behind a FH2 dimer (green/blue). B. Actin filament elongation by annealing of actin oligomers leaving behind a FH2 dimer (green/blue). Actin monomers or oligomers addition in a canonical actin helix conformation (helical twist of 167°) would be allowed at the formin-capped barbed ends within the open conformation. This would be most favorable when the leading FH2 domain is at a maximal distance from the helical axis, minimizing any steric clash with incoming actin subunits

**Supplementary text**

We discuss here how previously published data can lead to different estimates of the occupancy rate of the open state, depending on the hypotheses made for the theoretical analysis.

*Background*

In (Jegou et al. 2013), an increase in the mDia1-assisted elongation rate was measured as piconewton tensile forces were applied to the growing filaments. The formins were anchored to the coverslip surface, and the formins in interaction with the barbed ends thus experienced the pulling force applied to the filaments. For different profilin-actin concentrations (with a 3 µM excess of profilin) the elongation rate increased with force, reaching a plateau where elongation was roughly 2-fold faster than in the absence of force, and reaching half-maximum for a force of approximately 1 pN.

These experimental curves were fitted by a theoretical model, computed within the frame of the stair-stepping model, as follows.

In the absence of force, the elongation rate of the filament can be written as ***(equation 1)***:

$$v_{elong} = p_o \, k_{on} \, (C - C_c)$$

where $p_o$ the probability to be in the open conformation (i.e., the occupancy rate of the open state), $k_{on}$ is the on-rate constant for profilin-actin at the barbed end when the formin is in the open conformation, $C$ the profilin-actin concentration, and $C_c$ the critical concentration. Since, in most conditions, $C \gg C_c$ , we leave $C_c$ out of the following equations.

When a tensile force $F$ is applied to the formin, one can use transition state theory to compute how it changes the probability to be in the open state $p(F)$. Considering that the formin FH2 dimer is in rapid equilibirum between the open and the closed state as one FH2 domain moves back and forth between two positions separated by distance $\delta$, which is the actin monomer size, and assuming that $k_{on}$ is not affected by force, this leads to ***(equation 2)***:

$$v_{elong}(F) = \frac{k_{on} \, C}{1 + \frac{1-p_o}{p_o} e^{-F\delta/kT}}$$

there $T$ is temperature and $k$ is Boltzmann's constant.

Fitting the experimental curves for $v_{elong}(F)$ with this formula yielded $p_o = 0.56 \pm 0.06$ (and $k_{on} \approx 85 \ \mu\text{M}^{-1}\text{s}^{-1}$) (Jegou et al. 2013).

*Alternative analysis*

The theoretical curve (equation 2) was computed by considering that the force $F$ was applied to the mobile FH2 domain, at each step. An alternative would be to consider that only one FH2 domain is anchored to the surface, and thus that the force is exerted on this FH2 domain only. The on-rate would hence be affected only when this FH2 domain is the mobile, translocating domain. Therefore, in this alternative model, half of the subunits are added at the same rate as in the absence of force (following equation 1) while the other half of the subunits are added at the force-dependent rate (following equation 2), leading to ***(equation 3)***:

$$v_{elong}(F) = \frac{\frac{2\,p_o}{1+\ p_o} k_{on}\, C}{1 + \frac{1-\,p_o}{1+\,p_o} e^{-F\delta/kT}}$$

which has the same structure as equation 2. It can be rewritten as ***(equation 4)***:

$$v_{elong}(F) = \frac{k_{on}^{MOCK}\, C}{1 + \frac{1-p_o^{MOCK}}{p_o^{MOCK}} e^{-F\delta/kT}}$$

with $p_o^{MOCK} = \frac{1+p_o}{2}$ and $k_{on}^{MOCK} = \frac{2\,p_o}{1+\,p_o} k_{on}$.

Since equation 4 is basically the same as equation 3, it can fit the data equally, well with $p_o^{MOCK} = 0.56$ and $k_{on}^{MOCK} = 85\ \mu\text{M}^{-1}\text{s}^{-1}$.

This alternative model thus yields $p_o = 0.12 \pm 0.12$ (and $k_{on} \approx 397\ \mu\text{M}^{-1}\text{s}^{-1}$).
